# Utilizing CNNs for classification and uncertainty quantification for 15 families of European fly pollinators

**DOI:** 10.1101/2025.04.22.649975

**Authors:** Thomas Stark, Michael Wurm, Valentin Ştefan, Felicitas Wolf, Hannes Taubenböck, Tiffany M. Knight

## Abstract

Pollination is essential for maintaining biodiversity and ensuring food security, and in Europe it is primarily mediated by four insect orders (Coleoptera, Diptera, Hymenoptera, Lepidoptera). However, traditional monitoring methods are costly and time consuming. Although recent automation efforts have focused on butterflies and bees, flies, a diverse and ecologically important group of pollinators, have received comparatively little attention, likely due to the challenges posed by their subtle morphological differences. In this study, we investigate the application of Convolutional Neural Networks (CNNs) for classifying 15 European pollinating fly families and quantifying the associated classification uncertainty. Our dataset comprises a wide range of morphological and phylogenetic features, such as wing venation patterns and wing shapes. We evaluated the performance of three state-of-the-art CNN architectures, ResNet18, MobileNetV3, and EfficientNetB4, and demonstrate their effectiveness in accurately distinguishing fly families. In particular, EfficientNetB4 achieved an overall accuracy of up to 95.61%. Furthermore, cropping images to the bounding boxes of the Diptera not only improved classification accuracy but also increased prediction confidence, reducing misclassifications among families. This approach represents a significant advance in automated pollinator monitoring and has promising implications for both scientific research and practical applications.

## Introduction

Globally, insects are the most important animal pollinators, and therefore monitoring and understanding insect pollination is critical to biodiversity conservation and food security [1–3]. Current insect monitoring methods are expensive, labor-intensive, and slow, underscoring the need for efficient and automated approaches. Recent automation efforts have predominantly targeted butterflies and bees [4, 5]. However, fly pollinators provide substantial pollination services to wild and crop plants [6]. In Europe, 15 fly families are known to contribute to pollination, representing over 5000 species [7]. Classifying flies from images poses challenges due to morphological similarities across some families [8]. The limited availability of images for flies, compared to more charismatic taxa of pollinators, makes it difficult to create a comprehensive dataset, which is essential for successfully training Convolutional Neural Networks (CNNs) [9, 10].

Recent years have witnessed the widespread adoption of deep learning models for image classification, significantly advancing various fields, such as medicine [11], urban studies [12–14], and agricultural sciences [15, 16]. Furthermore, deep learning methods have increasingly become a key tool in the field of ecology and biodiversity, offering new avenues for research and conservation.

One of the most significant contributions of deep learning is in species identification and monitoring. The development of deep learning tools for the classification of arthropods initially focused on specific taxa (e.g., 16 species of mosquitoes) and the identification of museum species with uniform image backgrounds [17]. For example, images of arthropod wings taken in controlled settings, such as under a microscope, have been used to identify various groups of bees [18], butterflies [19, 20], and syrphid flies [21]. The use of CNNs for the identification of arthropods has expanded from a few taxa to multiple taxa, utilizing an increasing number of images. Examples include the identification of nine genera of tiger beetles with 380 images [22], eight groups of arthropods with nearly 20,000 images [23], and 36 species of bumblebee with nearly 90,000 images [24].

Using image classification techniques for pollinator identification often lacks insight into the confidence levels in its predictions. This is particularly relevant in challenging cases where image quality or species similarity can impact accuracy [17, 25]. Ensuring trustworthiness is critical and uncertainties play a crucial role in this case. This study delves into integrating uncertainty estimation methods within deep learning models to enhance predictive trustworthiness by explicitly addressing aleatoric and epistemic uncertainties.

The uncertainty estimation goes beyond traditional predictions, providing insight into the confidence and reliability of the model [26]. Incorporating these uncertainty estimation methods is crucial in real-world applications where decision-making about pollination abundance counts is based on accurate and reliable predictions. By transparently quantifying uncertainties, the models not only improve interpretability, but also provide a measure of confidence in their predictions [27]. This study outlines the methodologies used to capture and quantify uncertainties, contributing to the broader goal of establishing trust in deep learning models for image classification. One key research question is whether CNNs can effectively differentiate between the 15 families of pollinating flies, including a comparison of the impact of provided images versus cropping on classification accuracy.

This research focuses on the classification of a diverse range of Diptera families, employing a methodology that emphasizes both image refinement and robust confidence estimation. A key aspect involves comparing the performance of classification models when working with two distinct image datasets: The first dataset retains the original images sourced from the Global Biodiversity Information Facility (GBIF), while the second dataset relies on the same images but cropped to expert-defined bounding boxes that focus on the full specimen. To thoroughly evaluate these strategies, the approach compares three state-of-the-art CNN architectures ResNet18, MobileNetV3, and EfficientNetB4 across both image sets. Using a focus on the targeted areas, this approach aims to highlight defining features and reduce visual noise, ultimately guiding the classifier to more pertinent details.

To ensure that the classification framework remains transparent and interpretable, the research integrates uncertainty quantification methods. These techniques generate meaningful confidence values that reflect the level of certainty of the model, allowing researchers to better understand the quality and reliability of the predictions [28–30]. Such insights are particularly important for closely related Diptera families, where subtle differences in morphology may lead to misclassifications [8]. By examining groups with overlapping characteristics, specifically families such as Fannidae, Muscidae, and Tachinidae, the research attempts to refine classification boundaries and improve discriminative abilities.

## Materials and methods

### Describing 15 Diptera families

This research focuses on a set of 15 fly families selected not only for their distinctive morphological and phylogenetic traits, but also because they are known to provide pollination services throughout Europe [7]. A visual example fo each of the 15 families can be seen in Figure 1. These particular families embody a diverse array of species richness and ecological functions, providing a testbed for exploring classification challenges. The variation in the number of species across these families is notable; for example, while Sepsidae comprises around 48 known species in Europe, Tachinidae includes approximately 877 known species [7]. Such disparities in species richness can introduce significant variability in the visual characteristics of the specimens, influencing classification difficulty and the subsequent performance of CNN models.

**Fig 1.**
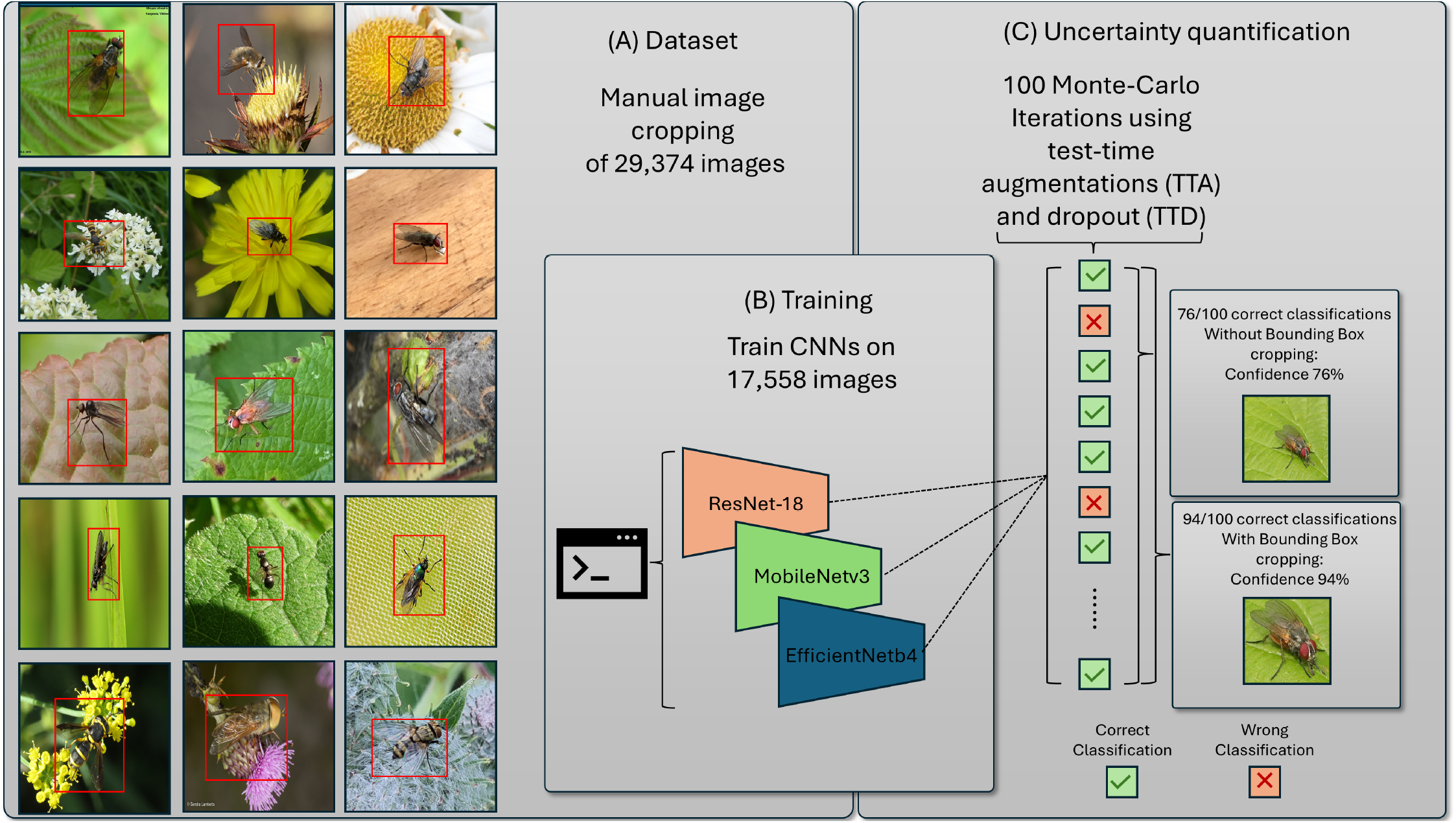
Uncertainty quantification approach. Methodology illustrating the creation of bounding boxes to crop Diptera images, training of CNNs on cropped and uncropped datasets, and confidence quantification using TTA and TTD with Monte Carlo iterations.

By investigating how morphological differences and evolutionary relationships shape the representation of these pollinating flies, this study aims to identify unique characteristics that can be effectively leveraged during the training and validation of classification models. Ultimately, understanding these nuances is critical to improving the accuracy, reliability, and interpretability of automated identification systems. In doing so, this research not only addresses the immediate task of classifying Diptera families, but also contributes to broader insights into how species diversity and phylogenetic relatedness affect computational taxonomy.

### Image acquisition

The images were obtained from GBIF with a focus on Europe, to maximize species richness. For families with abundant images, a random selection was applied to ensure a diverse dataset. Although it was not possible to automatically filter specifically for images of insects on flowers, efforts were made to remove images from museum collections, laboratory settings, fossils, duplicates, and other irrelevant sources or life stages. Due to varying levels of public interest, some species have a disproportionately large number of URLs available for image download, while others have significantly fewer resources. This results in a long-tail distribution of available images across the taxonomic spectrum of species and families. To address this imbalance, URLs were sampled as uniformly as possible across species within each family. Subsequently, a manual review was conducted to remove unwanted images, such as those depicting insect parts under magnification or misidentified specimens, thus ensuring a comprehensive and accurate dataset across species.

Expert-defined bounding boxes were created to enhance image accuracy by focusing on relevant features and reducing background noise. This approach is intended to improve the performance of CNN models in classifying the various families of pollinating flies.

### Data sampling

The dataset, comprising a total of 29,374 images, is partitioned into three distinct subsets, namely the training, validation, and testing datasets. To ensure a well-balanced and representative distribution across the 15 Diptera families, a stratified approach is employed. Each family contributes 60% of its images to the training set, allowing the models to learn from a wide and diverse range of examples. Subsequently, 20% of the images from each family are assigned to both the testing and validation sets. This strategic distribution guarantees that each Diptera family is adequately represented in all datasets, thus creating a comprehensive understanding of the nuances within the characteristics of each family. This data partitioning forms a critical aspect of model training and evaluation, facilitating robust and reliable performance assessments across diverse taxonomic groups.

### Selection of CNNs

In this study, we used three distinct CNNs to compare their performance on our dataset. MobileNetV3 Large, ResNet18, and EfficientNetB4. Each network was chosen based on specific attributes that align with our research goals.

1. MobileNetV3 Large [31] is designed to be both fast and efficient, making it particularly suitable for deployment in environments with limited computational resources. This model has approximately 5.4 million parameters, striking a balance between performance and efficiency. We selected MobileNetV3 Large to deliver high-speed inference without compromising accuracy significantly.
2. ResNet18 [32], a member of the Residual Networks family, is well-regarded for its robust performance across various scientific fields. With approximately 11.7 million parameters, ResNet18 is relatively lightweight, yet highly effective, thanks to its residual learning framework that eases the training of deep networks. This model was chosen for its proven efficacy in handling complex and diverse datasets.
3. EfficientNetB4 [33] is the largest of the three models in terms of parameters, with approximately 19 million parameters. It uses a compound scaling method that uniformly scales network dimensions, resulting in improved performance without the corresponding increase in computational cost. EfficientNetB4 is used to deliver high accuracy due to its larger size and advanced architectural design.

### Uncertainty quantification

Understanding and quantifying uncertainties in predictions are crucial for robust model deployment. We addressed two distinct forms of uncertainty, aleatoric and epistemic, through the application of Monte Carlo uncertainty approximations, whereby multiple classification outputs are averaged to derive confidence values.

Aleatoric uncertainty arises from inherent variability and randomness within the data itself [28]. Our model was designed to capture and quantify this type of uncertainty using test-time augmentations (TTA). This allows us to account for situations where the input data exhibit ambiguity or contain inherent noise. Aleatoric uncertainty might manifest itself in scenarios with subtle or ambiguous visual features.

We provide a random set of image augmentations during both the training phase and when the model is applied to the test dataset. This TTA approach ensures that the model encounters a diverse range of augmented inputs during training and testing, facilitating its ability to generalize and accurately assess uncertainty in real-world scenarios [34, 35].

Epistemic uncertainty, on the other hand, is rooted in the limitations of the model’s knowledge [28]. It reflects uncertainty arising from a lack of understanding or exposure to various data during training. We addressed epistemic uncertainty by incorporating test-time dropout (TTD). This helps the model recognize when faced with unfamiliar patterns not encountered during training, reducing the uncertainty associated with knowledge gaps [36].

Expanding on using Monte Carlo samples generated by TTD for uncertainty estimation [12, 34], our method extends the uncertainty estimation technique. We utilize Monte Carlo samples from TTA as well as from TTD. Our goal is to characterize the distribution, specifically the predictive posterior distribution of 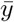. This is achieved by training the neural network as if it were a standard network, incorporating dropout layers after each layer with weight parameters and conducting *T* predictions.

Unlike the conventional classification scenario where a single prediction *y*^(*t*)^ is obtained, the combination of TTA and TTD techniques enables us to model a predictive distribution. This novel approach involves training the network as a typical neural network, but with slight modifications to the process.

### Experimental setup

In Figure 1, our methodological approach is illustrated in detail. Initially, bounding boxes are created for 29,374 images to crop them, ensuring that only the entire body of the Diptera is visible while removing any background as seen in Figure 1(A). This preprocessing step is crucial to focus the data set on the relevant features. Subsequently, three CNNs are trained using both the cropped and uncropped image datasets to assess the impact of background removal on classification performance as seen in Figure 1(B). Following this, we quantify the confidence of our predictions by approximating uncertainty estimates using TTA and TTD, with 100 Monte Carlo iterations providing robust statistical analysis. Finally, we compare the classification results, including confidence values, between the uncropped and cropped images, allowing a comprehensive evaluation of our methodological approach as seen in Figure 1(C).

The preprocessing pipeline for the images uses a z-score normalization for the images using the statistical parameters specific to the ImageNet dataset. This normalization ensures that the input images maintain consistency with the pre-training dataset, aligning their distribution with the expectations of the pre-trained model [32, 37].

Moreover, z-score normalization serves as a pivotal data preprocessing step due to its ability to standardize the input data, mitigating the impact of varying scales and intensities within the image data [38]. This is particularly crucial when working with diverse datasets that may exhibit significant variations in illumination conditions or imaging equipment. This improves the adaptability of the models to the unique features present in the Diptera image dataset, ultimately fostering their ability to capture and learn meaningful patterns during the training process.

Each CNN undergoes training for a consistent duration of 100 epochs. To ensure a robust evaluation, we present all metrics and results based on the model saved with the best validation score. In this context, the optimal model is determined by achieving the lowest validation score for the cross-entropy loss, employing label smoothing. This approach guarantees that the reported results reflect CNN’s performance at its peak during the training process, providing a comprehensive and accurate assessment of its capabilities.

In this study, we systematically evaluated the tradeoff between the per-epoch training speed of a CNN and the time required to achieve its best validation performance. Rapid training per epoch can reduce iteration times, but does not necessarily guarantee faster convergence to an optimal validation score. Our methods focused on identifying training configurations that balance these factors, thereby reducing the likelihood of overfitting and promoting better generalization to unseen data.

In addition, this evaluation was conducted with an emphasis on energy efficiency and environmental considerations, reflecting the principles of Green AI [39]. Specifically, we incorporated resource usage measurements and computational cost analyzes into our training protocols to ensure that performance gains did not come at the expense of excessive energy consumption [13]. Taking into account both training speed and environmental impact, this approach supports more responsible model development practices, aligning model optimization strategies with sustainability goals.

## Results

### Comparative analysis of CNNs

Table 3 compares three CNNs in terms of accuracy and efficiency, detailing their number of parameters, overall accuracy (OA), Kappa score, mean confidence, training epochs, time per epoch, test time for 100 Monte Carlo iterations, and whether the image was cropped to its bounding box.

**Table 1.** For each family the number of total images and the unique species and genus within each family within the dataset are shown.

| Family | Images | Species | Genera |
| --- | --- | --- | --- |
| Anthomyiidae | 1488 | 132 | 31 |
| Bombyliidae | 2239 | 151 | 40 |
| Calliphoridae | 1840 | 37 | 13 |
| Conopidae | 2356 | 57 | 10 |
| Empididae | 1632 | 144 | 14 |
| Fanniidae | 521 | 36 | 2 |
| Hybotidae | 1739 | 93 | 17 |
| Muscidae | 2163 | 212 | 41 |
| Sarcophagidae | 1346 | 77 | 22 |
| Scathophagidae | 2044 | 64 | 24 |
| Sepsidae | 2363 | 32 | 10 |
| Stratiomyidae | 2538 | 115 | 29 |
| Syrphidae | 2321 | 485 | 82 |
| Tabanidae | 2360 | 97 | 12 |
| Tachinidae | 2325 | 370 | 190 |

**Table 2.** Number of images in the Training (60%), Validation (20%), and Testing (20%) dataset.

| Family | Training | Validation | Testing |
| --- | --- | --- | --- |
| Anthomyiidae | 892 | 298 | 298 |
| Bombyliidae | 1343 | 448 | 448 |
| Calliphoridae | 1104 | 368 | 368 |
| Conopidae | 1413 | 471 | 472 |
| Empididae | 979 | 326 | 327 |
| Fanniidae | 312 | 104 | 105 |
| Hybotidae | 1043 | 348 | 348 |
| Muscidae | 1297 | 433 | 433 |
| Sarcophagidae | 807 | 269 | 270 |
| Scathophagidae | 1226 | 409 | 409 |
| Sepsidae | 1417 | 473 | 473 |
| Stratiomyidae | 1522 | 508 | 508 |
| Syrphidae | 1392 | 464 | 465 |
| Tabanidae | 1416 | 472 | 472 |
| Tachinidae | 1395 | 465 | 465 |
| **Total** | **17558** | **5856** | **5861** |

**Table 3.** Comparison of three CNNs in terms of accuracy and efficiency. The table includes the number of parameters, overall accuracy (OA), Kappa score, mean confidence, training epochs, time per epoch, test time for 100 Monte Carlo iterations, and if the image was cropped to its bounding box

| Model | Parameters | OA | Kappa | Mean Confidence | Epochs | Epoch Time | Test Time | Bounding Box Crop |
| --- | --- | --- | --- | --- | --- | --- | --- | --- |
| MobileNetV3 Large | 5.4m | 88.78% | .8887 | 78.49% | 47/100 | 00:05:16 | 01:53:27 | X |
| ResNet-18 | 11.3m | 88.58% | .8895 | 79.17% | 95/100 | 00:04:57 | 01:55:57 | X |
| EfficientNetB4 | 17.6m | 90.35% | .9044 | 79.27% | 46/100 | 00:11:34 | 02:22:32 | X |
| MobileNetV3 Large | 5.4m | 93.87% | .9401 | 85.85% | 94/100 | 00:05:23 | 01:46:30 | ✓ |
| ResNet-18 | 11.3m | 93.16% | .9384 | 83.98% | 54/100 | 00:04:46 | 01:49:16 | ✓ |
| EfficientNetB4 | 17.6m | 95.61% | .9590 | 87.38% | 70/100 | 00:11:23 | 02:29:24 | ✓ |

Overall, we see very high accuracies in the high 80% range for all model architectures and images or image sections, and in some cases well over 90%. Basically, we can therefore state that the 15 families can be classified very well. The overall accuracy (OA) for MobileNetV3 Large using the original images is 88.78%, while ResNet-18 and EfficientNetB4 achieve 88.58% and 90.35%, respectively.

The number of parameters for each model correlates with these results. MobileNetV3 Large, with 5.4 million parameters, shows lower OA compared to more complex models. ResNet-18, having 11.3 million parameters, performs similar to MobileNetV3 Large but slightly better. EfficientNetB4, which has the highest number of parameters at 17.6 million, consistently shows superior OA in both sets of experiments.

When images are cropped to their bounding box, the results generally show a significant improvement. For MobileNetV3 Large, the OA increases from 88.78% to 93.87%, which is an improvement of approximately 5.09. ResNet-18’s OA rises from 88.58% to 93.16%, an enhancement of about 4.58. EfficientNetB4 also benefits significantly, with its OA increasing from 90.35% to 95.61%, reflecting an improvement of approximately 5.26. In general, cropping images to their bounding box results in increased accuracies across the models, highlighting the importance of expert-defined bounding box image cropping in boosting model performance.

Examining the mean confidence, MobileNetV3 Large has a mean confidence of 78.49% for the original images and 85.85% when cropping the images in its bounding box. ResNet-18 records 79.17% and 83.98%, while EfficientNetB4 demonstrates 79.27% and 87.38%, respectively.

### Class confusion between Diptera families with and without Bounding Box Cropping

Figure 2 compares the confusion matrices for the EfficientNetB4 model, highlighting the classification performance across 15 Diptera families under two different conditions: (a) images cropped to their bounding box and (b) images not cropped.

**Fig 2.**
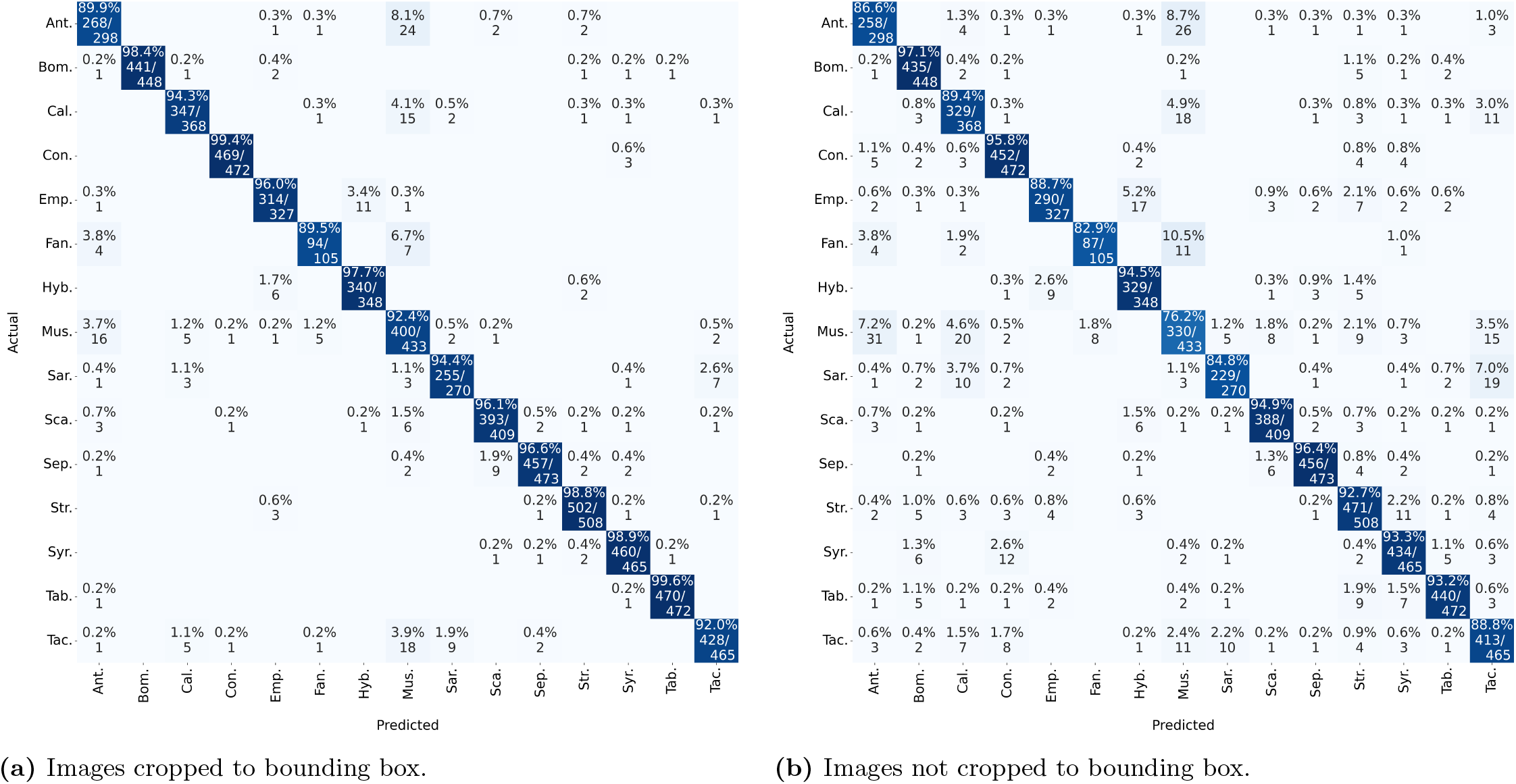
Confusion matrix comparing EfficientNetb4 results: (a) training with images cropped to the bounding box and (b) training with uncropped images.

From the matrices, it is evident that cropping images to their bounding box significantly enhances the model’s classification accuracy. This is reflected in higher true positive rates and reduced misclassification rates. For example, the Anthomyiidae family shows an accuracy of 89.9% with cropped images, compared to 86.6% with uncropped images. Similarly, for the Muscidae family, the accuracy is 92.4% when images are cropped, whereas it drops to 76.2% for uncropped images.

Moreover, the confusion within classes is noticeably higher when images are not cropped. This increased confusion is visible in the higher percentages of misclassification in several families. For example, in the Muscidae family, we observe that 7.2% of the samples are misclassified as Anthomyiidae, 4.6% as Calliphoridae and 3.5% as Tachinidae when the images are not cropped. In contrast, when the images are cropped, these misclassification rates drop significantly, showcasing more accurate predictions.

Similar patterns are observed in other families. For instance, the Sarcophagidae family, when images are uncropped, has misclassifications of 7.0% into Tachinidae and 3.7% into Calliphoridae. However, when cropped, the misclassification rates are significantly reduced, leading to more precise classification.

Overall, the results indicate that preprocessing images by cropping them to their bounding box substantially reduces confusion and improves the EfficientNetB4 model’s accuracy and reliability in classifying the 15 Diptera families.

### The uncertainty within the Diptera families

Figure 3 presents a boxplot illustrating the confidence values of the EfficientNetB4 model for all 15 Diptera families, comparing the results for the original images and the images cropped to their bounding box. This graph highlights the confidence levels for the correctly predicted images.

**Fig 3.**
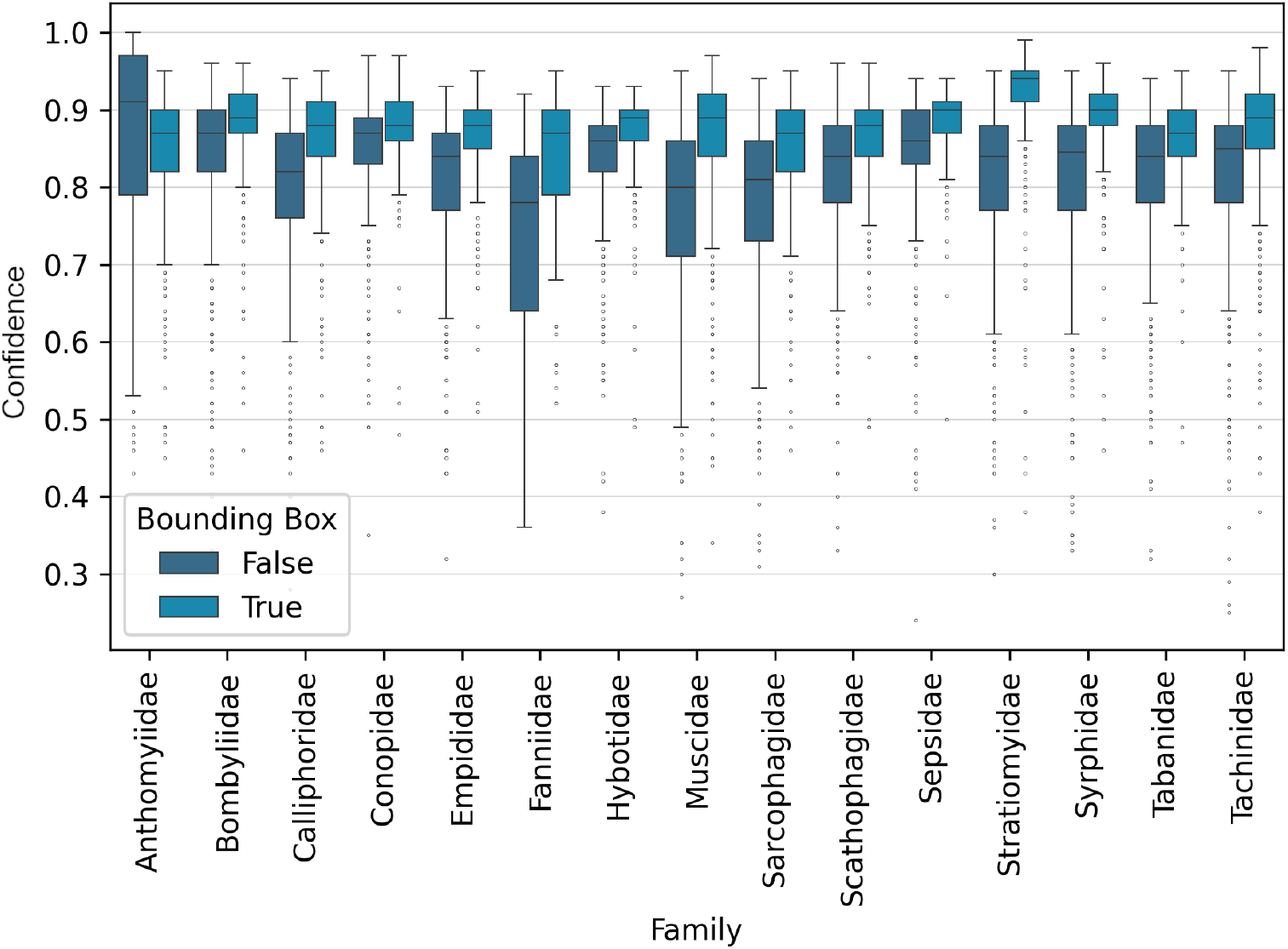
Confidence values for EfficientNetB4 for all 15 Diptera families for correct classification results with and without cropping the its Boundinx Boxes.

We sorted the families by the magnitude of improvement in their confidence scores, that is, the difference between the cropped image confidence and the original image confidence. Families with an increase of 6% or more are categorized as having large improvements and are listed first in descending order, while those with less than 6% improvement are grouped as small improvements, also in descending order.

In the group of large improvements, Stratiomyidae experiences the largest increase, with confidence increasing from 84% to 94% (a 10% improvement). Muscidae and Fanniidae follow with significant increases of 9% each, with Muscidae improving from 80% to 89% and Fanniidae from 78% to 87%. Both Calliphoridae and Sarcophagidae show a 6% improvement, with Calliphoridae moving from 82% to 88% and Sarcophagidae from 81% to 87%.

In the small improvements group, Syrphidae’s confidence increases from 85% to 90% (a 5% improvement). Sepsidae improve from 86% to 90% (a 4% increase), as do Empididae and Scathophagidae, each increasing from 84% to 88% (a 4% improvement). Tachinidae also shows a 4% increase, moving from 85% to 89%. Hybotidae experiences a 3% improvement, going from 86% to 89%, and Tabanidae likewise improves by 3%, rising from 84% to 87%. Bombyliidae sees a modest 2% increase from 87% to 89%, while Conopidae remains unchanged at 88%.

Anthomyiidae is the only family that shows a decrease in median confidence, falling from 91% to 87%.

### Visual results for all 15 families

The visual results of the analysis of the 15 Diptera families highlight significant differences between the two training approaches using the best performing model EfficientNetB4. Specifically, we compared the performance when the model was trained on images cropped to the Diptera bounding box versus images that were not cropped. Figure 4 presents two images from the test dataset for each Diptera family, highlighting both the highest and lowest confidence differences between the images cropped to its bounding box and the uncropped ones. Moreover, these images were automatically selected rather than hand-picked, ensuring an unbiased and representative illustration of the model’s performance. Generally, as illustrated in Figure 3, the model trained on cropped images consistently achieves better results.

**Fig 4.**
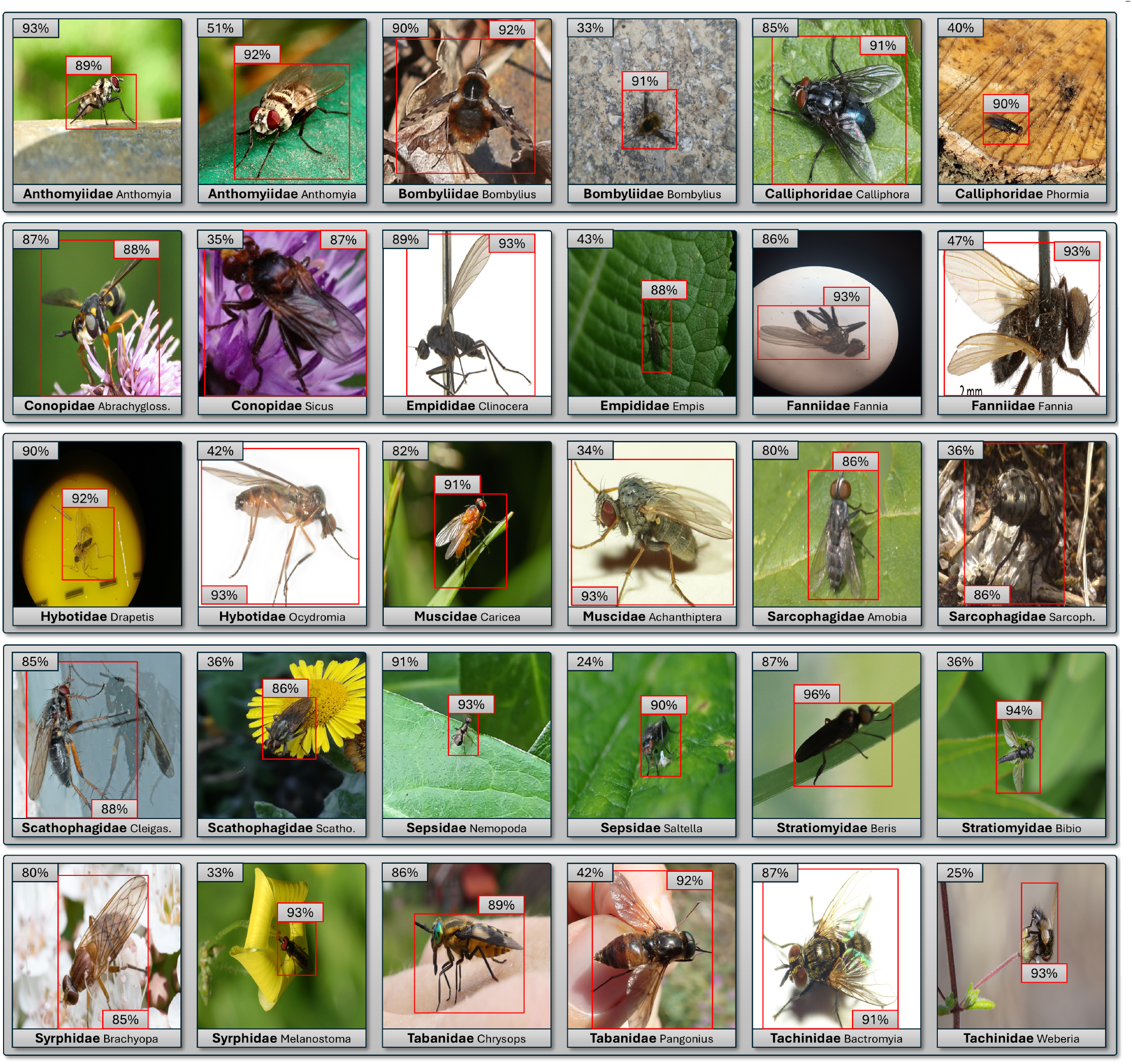
Comparison of EfficientNetB4 confidence values on 15 Diptera families. The figure highlights confidence differences between models trained on images cropped to the Diptera’s bounding box (red bounding box) versus those using (full) images. For each Diptera family, two examples are shown: one representing the smallest difference and one representing the largest difference in confidence values of the test dataset.

## Discussion

Figure 4 presents a comprehensive comparison of models trained with and without cropping to the Diptera’s bounding box across the 15 families. For enhanced visual clarity, the species for each family are displayed solely in this figure. Note that this information was not used in the classification process but is provided only to offer additional context for interpreting the results.

Creating bounding boxes manually is an extremely time-consuming process. For the 29,374 images used in this study, bounding boxes had to be created by experts, following a consistent workflow to ensure that each specimen is cropped uniformly. This task requires not only significant time but also specialized expertise to accurately delineate the objects of interest. The labor-intensive nature of this manual approach underscores the potential benefits of automation. It would be very interesting to explore the use of object detection algorithms to automatically crop images similar to the methods used in [40] and [23]. Moreover, comparing the performance of models trained on manually cropped images with those cropped automatically could provide valuable insights into both the feasibility and efficiency of automated cropping.

In most cases, cropping improves performance by focusing the model on the relevant features of the Diptera, thereby reducing background noise. This improvement is likely due to the model’s increased focus on Diptera features, which minimizes background noise and facilitates more effective feature extraction, which ultimately leads to greater overall precision [41]. For example, within the Calliphoridae family, the Phormia specimen occupies only a small fraction of the full image, resulting in a dramatic increase in the predicted confidence, from 40% without cropping to 90% with cropping. In contrast, for Calliphoridae Calliphora, where the diptera fills almost the entire image, the confidence values are much closer (91% with cropping vs. 85% without cropping). This trend of improved performance with cropping is evident in several families.

However, there are some notable outliers. In cases such as Conopidae Sicus, the second example of Fanniidae Fannia, Hybotidae Ocydromia, Muscidae Achanthiptera, and Tabanidae Pangonius, the bounding box is only slightly smaller than the original image, yet the differences between the cropped and non-cropped models are pronounced. In many of these cases, the uniform laboratory environment in the background could significantly influence the model’s training. In addition, challenging scenarios are observed in Sarcophagidae Sarcophaga, where the Diptera closely resembles the background, and in Scathophagidae Cleigastra, where the specimen is imaged on a highly reflective surface, both cases that test the model’s adaptability under varying conditions.

### Challenges in Diptera classification

Distinguishing between these 15 Diptera families from images is a challenging task, because many families share numerous morphological features, and certain distinguishing characteristics might not be visible in images. Figure 5 illustrates these challenges by comparing the results of three morphologically similar Diptera families. In such scenarios, even a trained expert struggles to confidently separate closely related groups by manual-visual interpretation, leading to potential misclassification when relying on visual cues alone. Despite these challenges, our models demonstrate robust performance. However, overfitting remains a concern in many deep learning algorithms. Models may exploit subtle cues that are not directly linked to the underlying taxonomy, and dataset sampling biases can also contribute to overfitting. This possibility underscores the critical need for our uncertainty estimation approach, which integrates extensive image augmentations and dropout to mitigate these effects.

**Fig 5.**
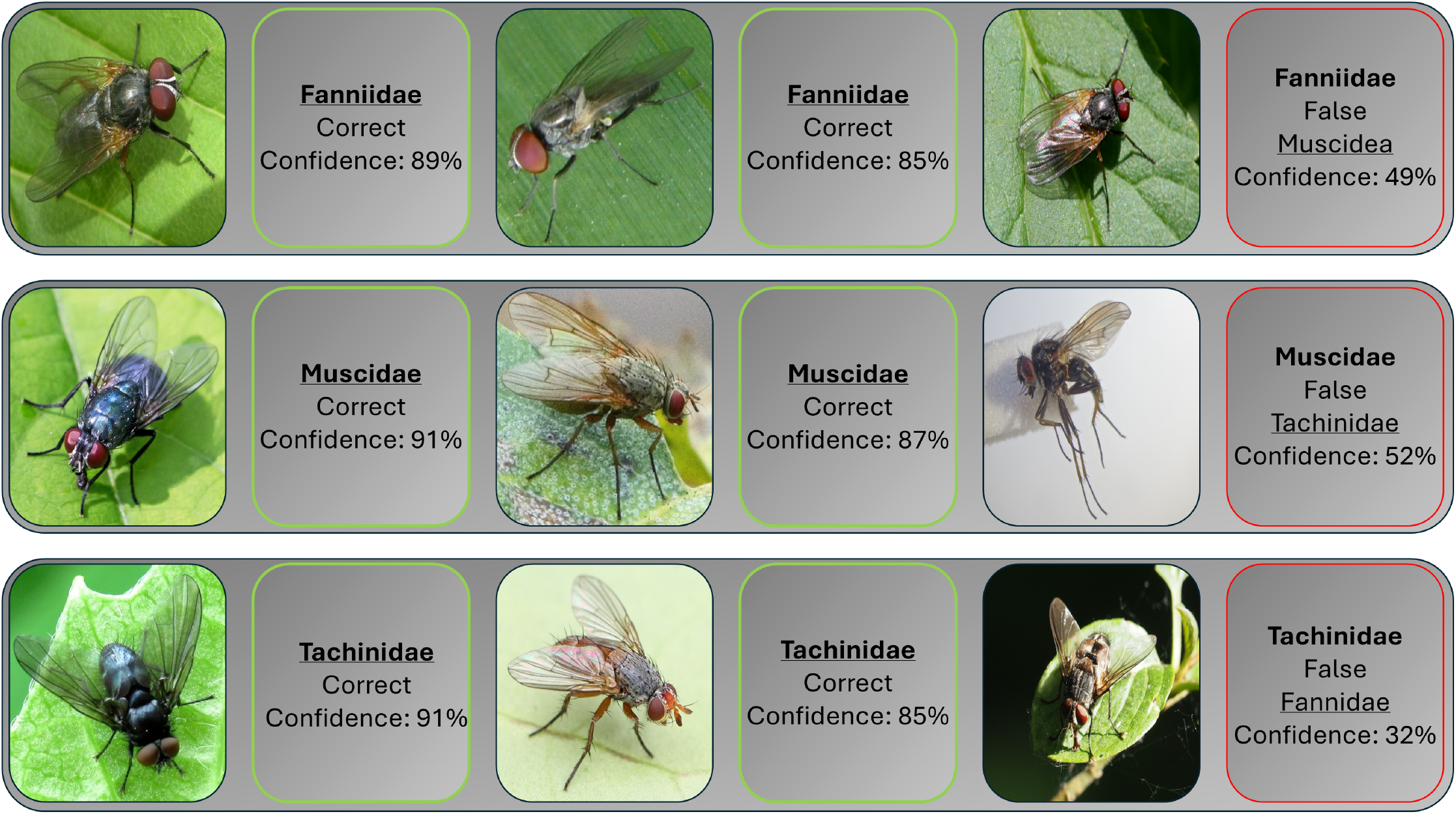
Confidence values for three Diptera families with morphological similarities. Bold family names are the labels (ground truth), while underlined family names are the predictions. Confidence scores are based on EfficientNetB4 with images cropped to their bounding boxes.

By incorporating 100 Monte Carlo iterations, we gain a quantitative window into the confidence scores of the model, enabling us to confirm that the impressive precision is neither coincidental nor artificially inflated. Instead, these analyzes demonstrate that the model’s capabilities extend beyond pattern matching and that it genuinely understands the underlying structure of the data. In fact, this examination reveals that the model can consistently identify subtle class distinctions, even in fuzzy feature spaces where related classes share overlapping traits. The combination of strong performance and transparent uncertainty quantification offers both confidence and credibility.

As observed in [40], cropping images to the insect’s bounding box has a significant effect on enhancing model performance. This improvement is expected since the models do not need to process excessive background information, which can be distracting. Distracting backgrounds can have a substantial impact on model accuracy.

The results of our study highlight the effectiveness of different CNN architectures in classifying Diptera families. Although earlier generation models such as ResNet-18 provide acceptable accuracy, more recent models, such as EfficientNet-B4, demonstrate better performance, offering higher confidence and fewer misclassifications. For scenarios where speed and portability are critical, lightweight models such as MobileNetv3 deliver strong results, enabling on-device classification without significantly compromising accuracy. In this way, we see a clear progression in model capabilities, from established architectures to cutting-edge networks, each offering specific advantages that can be leveraged according to the demands of the task.

### Confusion between Diptera families

Four families were most often confused with each other: Anthomyiidae, Fanniidae, Muscidae, and Tachinidae, all of which belong to the same superfamily, Muscoidea. Their morphological similarities often complicate accurate identification. In particular, wing venation patterns serve as a critical feature to distinguish these families, but it is challenging to capture clear and properly angled images to visualize the entire wing. Despite these difficulties, it is rewarding to see that the models perform as well as they did.

To illustrate this performance, we present confidence values for a subset of these morphologically similar Diptera families seen in Figure 5. Family names in bold indicate the true labels, while underlined family names represent the model’s predictions. These confidence scores, obtained using an EfficientNetB4 model with bounding box cropped images, underscore the model’s capability to differentiate among closely related taxa.

From the confusion matrix in Figure 2.a, we observe that of the 105 Fannidae images in our test dataset, seven are misclassified as Muscidae. One such misclassification is shown in Figure 5, where EfficientNet exhibits low confidence (49%) in its classification. In contrast, correctly classified Fannidae images have higher confidence scores of 85% and 89%.

For the Muscidae family, five of 433 test images are misclassified as Fannidae and two as Tachinidae. A misclassification of Tachinidae is depicted in Figure 5, with a low confidence of 52%. Correct classifications for Muscidae have higher confidence scores of 87% and 91%.

In the Tachinidae family, one out of 465 test images is misclassified as Fannidae, and 18 as Muscidae. The single misclassification as Fannidae is shown in Figure 5, with a very low confidence score of 32%. The correct classifications of Tachinidae have confidence scores of 85% and 91%. Thus, in general, for the purposes of ecological studies, when confidence scores are 85% or higher, it is reasonable to identify the fly in the image to the family taxonomic level. However, when the confidence level is less than 50%, it is advisable to keep the identification at the order level (Diptera).

The few misclassifications observed are informative as well. They highlight the areas where the model can be further refined, and they provide insight into the limits of current image recognition technology in dealing with highly similar morphological features. This approach is a significant step forward in the field of automated species identification and has the potential to contribute meaningfully to various scientific and practical applications.

The next step in this line of research is to delve deeper into the taxonomic hierarchy. The ability to classify pollinating insects at the species levels automatically from images would be highly beneficial to ecological monitoring [24]. This progression will test the limits of our current methodologies and may necessitate the development of new approaches. Although our existing models have proven effective at higher taxonomic levels, the increased specificity required at the species levels could demand more sophisticated architectures, such as transformers, as well as larger and more well sampled datasets. We anticipate that this new challenge will be complex, yet we remain optimistic about our ability to overcome potential obstacles. Adapting and refining our methods to achieve fine-scale classification will not only advance our understanding of pollinator diversity but also contribute to broader ecological and conservation efforts.

## Conclusion

In conclusion, our study demonstrates that automated pollinator monitoring using CNNs can overcome many of the limitations of traditional methods, offering a cost-effective and efficient alternative resulting in high classification accuracies. By focusing on 15 European pollinating fly families, a group that has traditionally been challenging due to subtle morphological differences, we showed that CNN architectures such as ResNet18, MobileNetV3, and especially EfficientNetB4, can achieve high classification accuracy (from 88.58% to 95.61%). A key finding was that cropping images to the Diptera’s bounding boxes not only improved accuracy but also enhanced prediction certainty, effectively reducing misclassifications among families.

Visual analyses further corroborate these results. In most cases, especially when the fly occupies only a small portion of the image, cropping significantly increased the predicted confidence, thereby reinforcing the benefits of focusing on the relevant features. Although some outliers were observed, possibly due to uniform lab environments or challenging imaging conditions, the overall trend supports the use of bounding-box cropping as a robust method for improving model performance.

This work not only advances the field of automated pollinator monitoring, but also provides a foundation for future applications in ecological research and practical conservation efforts, ensuring that critical pollination services are better understood and protected.

## Data and code availability

The code used to collect and process the data is publicly available on our GitHub repository. You can access both the data collection code and the data processing code here stark-t/PAI diptera.

## Author contributions

**Conceptualization:** Thomas Stark, Michael Wurm, Valentin Stefan, Hannes Taubenboeck, Tiffany Knight.

**Data curation:** Thomas Stark, Valentin Stefan, Felicitas Wolf.

**Formal analysis:** Thomas Stark, Michael Wurm, Valentin Stefan, Tiffany Knight.

**Investigation:** Thomas Stark, Michael Wurm, Valentin Stefan, Tiffany Knight.

**Methodology:** Thomas Stark.

**Project administration:** Thomas Stark, Michael Wurm, Valentin Stefan, Hannes Taubenboeck, Tiffany Knight.

**Resources:** Thomas Stark, Michael Wurm, Valentin Stefan, Hannes Taubenboeck, Tiffany Knight.

**Software:** Thomas Stark, Valentin Stefan, Felicitas Wold.

**Supervision:** Michael Wurm, Hannes Taubenboeck, Tiffany Knight.

**Validation:** Thomas Stark, Michael Wurm, Valentin Stefan, Hannes Taubenboeck, Tiffany Knight.

**Visualization:** Thomas Stark.

**Writing – original draft:** Thomas Stark, Tiffany Knight.

**Writing – review & editing:** Michael Wurm, Valentin Stefan, Felicitas Wolf, Hannes Taubenboeck.

## References

1. Klein AM, Vaissière BE, Cane JH, Steffan-Dewenter I, Cunningham SA, Kremen C, et al. Importance of pollinators in changing landscapes for world crops. Proceedings of the royal society B: biological sciences. 2007;274(1608):303–313. doi:10.1098/rspb.2006.3721.

2. Hallmann CA, Sorg M, Jongejans E, Siepel H, Hofland N, Schwan H, et al. More than 75 percent decline over 27 years in total flying insect biomass in protected areas. PLOS ONE. 2017;12(10):1–21. doi:10.1371/journal.pone.0185809.

3. Sánchez-Bayo F, Wyckhuys KAG. Worldwide decline of the entomofauna: A review of its drivers. Biological Conservation. 2019;232:8–27. doi:10.1016/j.biocon.2019.01.020.

4. Alves TS, Pinto MA, Ventura P, Neves CJ, Biron DG, Junior AC, et al. Automatic detection and classification of honey bee comb cells using deep learning. Computers and Electronics in Agriculture. 2020;170:105244. doi:10.1016/j.compag.2020.105244.

5. Cunha C, Narotamo H, Monteiro A, Silveira M. Detection and measurement of butterfly eyespot and spot patterns using convolutional neural networks. PLOS ONE. 2023;18(2):1–15. doi:10.1371/journal.pone.0280998.

6. Rader R, Bartomeus I, Garibaldi LA, Garratt MPD, Howlett BG, Winfree R, et al. Non-bee insects are important contributors to global crop pollination. Proceedings of the National Academy of Sciences. 2016;113(1):146–151. doi:10.1073/pnas.1517092112.

7. Pape T, Beuk P, Pont AC, Shatalkin AI, Ozerov AL, Woźnica AJ, et al. Fauna Europaea: Diptera – Brachycera. Biodiversity Data Journal. 2015;3:e4187. doi:10.3897/BDJ.3.e4187.

8. Yeates DK, Wiegmann BM. Phylogeny and evolution of Diptera: recent insights and new perspectives. The evolutionary biology of flies. 2005; p. 14–44.

9. Boakes EH, McGowan PJK, Fuller RA, Chang-qing D, Clark NE, O’Connor K, et al. Distorted Views of Biodiversity: Spatial and Temporal Bias in Species Occurrence Data. PLOS Biology. 2010;8(6):1–11. doi:10.1371/journal.pbio.1000385.

10. Troudet J, Grandcolas P, Blin A, Vignes-Lebbe R, Legendre F. Taxonomic bias in biodiversity data and societal preferences. Scientific reports. 2017;7(1):9132.

11. Litjens G, Kooi T, Bejnordi BE, Setio AAA, Ciompi F, Ghafoorian M, et al. A survey on deep learning in medical image analysis. Medical Image Analysis. 2017;42:60–88. doi:10.1016/j.media.2017.07.005.

12. Stark T, Wurm M, Zhu XX, Taubenböck H. Quantifying Uncertainty in Slum Detection: Advancing Transfer Learning With Limited Data in Noisy Urban Environments. IEEE Journal of Selected Topics in Applied Earth Observations and Remote Sensing. 2024;17:4552–4565. doi:10.1109/JSTARS.2024.3359636.

13. Stiller D, Stark T, Strobl V, Leupold M, Wurm M, Taubenböck H. Efficiency of CNNs for Building Extraction: Comparative Analysis of Performance and Time. In: 2023 Joint Urban Remote Sensing Event (JURSE); 2023. p. 1–4.

14. Wurm M, Stark T, Zhu XX, Weigand M, Taubenböck H. Semantic segmentation of slums in satellite images using transfer learning on fully convolutional neural networks. ISPRS Journal of Photogrammetry and Remote Sensing. 2019;150:59–69. doi:10.1016/j.isprsjprs.2019.02.006.

15. Rußwurm M, Courty N, Emonet R, Lefèvre S, Tuia D, Tavenard R. End-to-end learned early classification of time series for in-season crop type mapping. ISPRS Journal of Photogrammetry and Remote Sensing. 2023;196:445–456. doi:10.1016/j.isprsjprs.2022.12.016.

16. Kamilaris A, Prenafeta-Boldú FX. A review of the use of convolutional neural networks in agriculture. The Journal of Agricultural Science. 2018;156(3):312–322. doi:10.1017/S0021859618000436.

17. Wäldchen J, Mäder P. Machine learning for image based species identification. Methods in Ecology and Evolution. 2018;9(11):2216–2225. doi:10.1111/2041-210X.13075.

18. Buschbacher K, Ahrens D, Espeland M, Steinhage V. Image-based species identification of wild bees using convolutional neural networks. Ecological Informatics. 2020;55:101017. doi:10.1016/j.ecoinf.2019.101017.

19. Kaya Y, Kayci L, Uyar M. Automatic identification of butterfly species based on local binary patterns and artificial neural network. Applied Soft Computing. 2015;28:132–137. doi:10.1016/j.asoc.2014.11.046.

20. Almryad AS, Kutucu H. Automatic identification for field butterflies by convolutional neural networks. Engineering Science and Technology, an International Journal. 2020;23(1):189–195. doi:10.1016/j.jestch.2020.01.006.

21. Mielczarek E, Tofilski A. Semiautomated identification of a large number of hoverfly (Diptera: Syrphidae) species based on wing measurements. Oriental insects, 52(3), 245–258.

22. Abeywardhana DL, Dangalle CD, Nugaliyadde A, Mallawarachchi Y. Deep learning approach to classify Tiger beetles of Sri Lanka. Ecological Informatics. 2021;62:101286. doi:10.1016/j.ecoinf.2021.101286.

23. Stark T, Ştefan V, Wurm M, Spanier R, Taubenböck H, Knight TM. YOLO object detection models can locate and classify broad groups of flower-visiting arthropods in images. Scientific Reports. 2023;13(1):16364.

24. Spiesman BJ, Gratton C, Hatfield RG, Hsu WH, Jepsen S, McCornack B, et al. Assessing the potential for deep learning and computer vision to identify bumble bee species from images. Scientific Reports. 2021;11(1):7580. doi:10.1038/s41598-021-87210-1.

25. Valan M, Makonyi K, Maki A, Vondráček D, Ronquist F. Automated Taxonomic Identification of Insects with Expert-Level Accuracy Using Effective Feature Transfer from Convolutional Networks. Systematic Biology. 2019;68(6):876–895. doi:10.1093/sysbio/syz014.

26. Abdar M, Pourpanah F, Hussain S, Rezazadegan D, Liu L, Ghavamzadeh M, et al. A review of uncertainty quantification in deep learning: Techniques, applications and challenges. Information Fusion. 2021;76:243–297. doi:10.1016/j.inffus.2021.05.008.

27. Pham N, Fomel S. Uncertainty and interpretability analysis of encoder-decoder architecture for channel detection. GEOPHYSICS. 2021;86(4):O49–O58. doi:10.1190/geo2020-0409.1.

28. Kendall A, Gal Y. What Uncertainties Do We Need in Bayesian Deep Learning for Computer Vision? In: Guyon I, Luxburg UV, Bengio S, Wallach H, Fergus R, Vishwanathan S, et al., editors. Advances in Neural Information Processing Systems. vol. 30. Curran Associates, Inc.; 2017. Available from: https://proceedings.neurips.cc/paper_files/paper/2017/file/2650d6089a6d640c5e85b2b88265dc2b-Paper.pdf.

29. Valdenegro-Toro M. I Find Your Lack of Uncertainty in Computer Vision Disturbing. In: Proceedings of the IEEE/CVF Conference on Computer Vision and Pattern Recognition (CVPR) Workshops; 2021. p. 1263–1272.

30. Huang L, Lala S, Jha NK. CONFINE: Conformal Prediction for Interpretable Neural Networks; 2024. Available from: https://arxiv.org/abs/2406.00539.

31. Howard A, Sandler M, Chu G, Chen LC, Chen B, Tan M, et al. Searching for MobileNetV3. In: Proceedings of the IEEE/CVF International Conference on Computer Vision (ICCV); 2019.

32. He K, Zhang X, Ren S, Sun J. Deep Residual Learning for Image Recognition; 2015. Available from: https://arxiv.org/abs/1512.03385.

33. Tan M, Le QV. EfficientNet: Rethinking Model Scaling for Convolutional Neural Networks; 2020. Available from: https://arxiv.org/abs/1905.11946.

34. Wang G, Li W, Aertsen M, Deprest J, Ourselin S, Vercauteren T. Aleatoric uncertainty estimation with test-time augmentation for medical image segmentation with convolutional neural networks. Neurocomputing. 2019;338:34–45. doi:10.1016/j.neucom.2019.01.103.

35. MacIsaac J, Newson S, Ashton-Butt A, Pearce H, Milner B. Improving acoustic species identification using data augmentation within a deep learning framework. Ecological Informatics. 2024;83:102851. doi:10.1016/j.ecoinf.2024.102851.

36. Nappa A, Quartulli M, Azpiroz I, Marchi S, Guidotti D, Staiano M, et al. Probabilistic Bayesian Neural Networks for olive phenology prediction in precision agriculture. Ecological Informatics. 2024;82:102723. doi:10.1016/j.ecoinf.2024.102723.

37. Simonyan K. Very deep convolutional networks for large-scale image recognition. arXiv preprint arXiv:14091556. 2014;.

38. Kamnitsas K, Ledig C, Newcombe VF, Simpson JP, Kane AD, Menon DK, et al. Efficient multi-scale 3D CNN with fully connected CRF for accurate brain lesion segmentation. Medical image analysis. 2017;36:61–78.

39. Desislavov R, Martínez-Plumed F, Hernández-Orallo J. Trends in AI inference energy consumption: Beyond the performance-vs-parameter laws of deep learning. Sustainable Computing: Informatics and Systems. 2023;38:100857. doi:10.1016/j.suscom.2023.100857.

40. Sittinger M, Uhler J, Pink M, Herz A. Insect detect: An open-source DIY camera trap for automated insect monitoring. PLOS ONE. 2024;19(4):1–28. doi:10.1371/journal.pone.0295474.

41. Choton JC, Margapuri V, Grijalva I, Spiesman B, Hsu WH. Self-supervised Component Segmentation To Improve Object Detection and Classification For Bumblebee Identification. bioRxiv. 2025; p. 2025–03.

